# Terrestriality and bacterial transfer: A comparative study of gut microbiomes in sympatric Malagasy mammals

**DOI:** 10.1101/293282

**Authors:** Amanda C. Perofsky, Rebecca J. Lewis, Lauren Ancel Meyers

**Author notes:** **Correspondence**: AC Perofsky, Department of Integrative Biology, The University of Texas at Austin, 2415 Speedway #C0930, Austin, Texas, 78712, USA.

## Abstract

The gut microbiomes of mammals appear to mirror their hosts’ phylogeny, suggesting a shared history of co-speciation. Yet, much of this evidence stems from comparative studies of distinct wild or captive populations that lack data for disentangling the relative influences of shared phylogeny and environment. Here, we present phylogenetic and multivariate analyses of gut microbiomes from six sympatric (*i.e.*, co-occurring) mammal species inhabiting a 1-km^2^ area in western Madagascar—three lemur and three non-primate species—that consider genetic, dietary, and ecological predictors of microbiome functionality and composition. Host evolutionary history, indeed, appears to drive gut microbial patterns among distantly related species. However, we also find that diet—reliance on leaves versus fruit—is the best predictor of microbiome similarity among closely related lemur species, and that host substrate preference—ground versus tree— constrains horizontal transmission via incidental contact with feces, with arboreal species harboring far more distinct communities than those of their terrestrial and semi-terrestrial counterparts.

## Introduction

The gastrointestinal tracts of mammals are complex ecosystems harboring large and diverse populations of bacteria that are essential for digestion, development, metabolism, behavior, immune function, and protection from pathogens (1–6). These microbial communities are potentially shaped by diverse *host* factors – heritable (*e.g.*, genetics or evolutionary history), environmental (*e.g.*, geography, diet), as well as behavioral (*i.e.*, social contact patterns) (7). Hypothetically, if gut bacteria colonize hosts strictly via maternal inheritance (*i.e.*, vertical transmission) or intraspecific transmission, then gut microbiota should stably co-diversify with their hosts and thus mirror their evolutionary relationships (8,9). Conversely, if they are horizontally transmitted among distantly related hosts, via shared food sources or habitat, then animals with overlapping diets and environmental exposures should harbor similar microbiomes, regardless of their phylogenetic relationships (9). However, dietary and habitat preferences can obscure the phylogenetic signal within host-associated microbial communities, thus confounding attempts to estimate the relative influences of vertical, horizontal, and environmental transmission on mammalian gut microbiome composition (9).

Several studies have estimated the relative influences of host phylogeny, genetics, diet, and environment on both interspecific and intraspecific microbiome diversity in mammals. Several studies of zoo animals reared on artificial diets have claimed that diet and shared environment have greater impacts on gut microbiome composition than endogenous factors (10–14). Other studies have focused on closely related mammal species, and generally found that host species and their gut microbes exhibit concordant phylogenies (8,15–18), with the gut microbiomes of different host species forming distinct clusters (18–20). Within single host species, some studies of humans and other primates have suggested that social contact patterns are the primary predictor of gut microbiome composition (21–25), whereas others have found that host genetics exert a strong influence (26,27).

Most of these comparative microbiome surveys have sampled geographically and ecologically distinct captive or wild populations (but see 14,27,28). However, captivity alters microbiome composition in mammals (12,30–32), and, when animals are compared across geographic regions, host phylogenetic differences may be confounded by differences in local microbial taxa (15,33). Thus, many of these prior studies did not have sufficient data for resolving the relative influences of various heritable and environmental factors.

Madagascar is home to a unique and threatened constellation of mammalian fauna, with high levels of endemism and broad species diversity across remarkably few taxonomic groups (34,35). This phenomena has been attributed to Madagascar’s long isolation from other continents, which predates the evolution of most recent mammals, and rare “sweepstake” migration events over the past 65 million years (36,37). All non-flying mammals fall into either one of four endemic orders—lemurs, tenrecs, carnivorans, or rodents—or are recent human introductions—African bush pigs (38), cattle, cats, or dogs (39). Recent studies have shown that social contacts and diet influence the gut microbiomes of wild lemurs (24,25,40), but none have yet considered the microbiomes of co-occurring mammalian species.

Here, we present an analysis of microbiome diversity across six sympatric (*i.e.*, geographically co-occurring) mammalian populations inhabiting a 1-km^2^ area in western Madagascar. The focal species include both folivorous and frugivorous lemurs (Verreaux’s sifaka, *Propithecus verreauxi*; red-tailed sportive lemur, *Lepilemur ruficaudatus*; red-fronted brown lemur, *Eulemur rufifrons*), a viverrid that is the largest extant carnivore in Madagascar (fossa, *Cryptoprocta ferox*), and two human-introduced artiodactyls (even-toed ungulates) – one wild (African bush pig, *Potamochoerus larvatus*) and the other domesticated (zebu cattle, *Bos t. indicus*). We utilize multivariate modeling and phylogenetic approaches to disentangle the relative contributions of host environment, diet, and evolutionary history on gut microbial community structure. We estimate microbiome diversity, composition, and functional potential using 16S rRNA sequences from fecal samples of 61 individual animals as well as published surveys and show that host phylogeny predicts compositional diversity on a broad scale, while host diet appears to exert a greater influence among recently diverged primate hosts. Furthermore, we find evidence of greater horizontal transmission between distantly related *terrestrial* mammals, relative to closely related *arboreal* species, thus suggesting that substrate preference (*i.e.*, the primary surface upon which animals locomote) shapes microbiome composition.

## Materials and Methods

### Fecal sample collection

We collected fresh fecal samples from wild populations of Verreaux’s sifaka (*Propithecus verreauxi*), red-tailed sportive lemurs (*Lepilemur ruficaudatus*), red-fronted brown lemurs (*Eulemur rufifrons*), fossa (*Cryptoprocta ferox*), and African bush pigs (*Potamochoerus larvatus*) within a 1-km^2^ area at Ankoatsifaka Research Station (20°47.69′S, 44°9.88′E; Figs. 1C and S1) in Kirindy Mitea UNESCO Biosphere Reserve in western Madagascar. During a two-month span in the dry season (6 June 2012 – 1 August 2012), we collected sifaka (N = 14 individuals), sportive lemur (N = 14), and brown lemur samples (N = 11) during individual animal follows, after individuals were observed defecating, and fossa (N = 6) and bush pig samples (N = 8) opportunistically within the research site. We collected cattle samples (N = 8) along a 20 km stretch of dirt road that traverses Kirindy Mitea. We recorded the GPS location of each sample at the time of collection (Table S1) and preserved samples in RNAlater® (ThermoFisher, Waltham, MA, USA) at ambient temperature until their arrival at the University of Texas at Austin in August 2012. We subsequently stored samples at −80°C until further processing.

**Figure 1.**
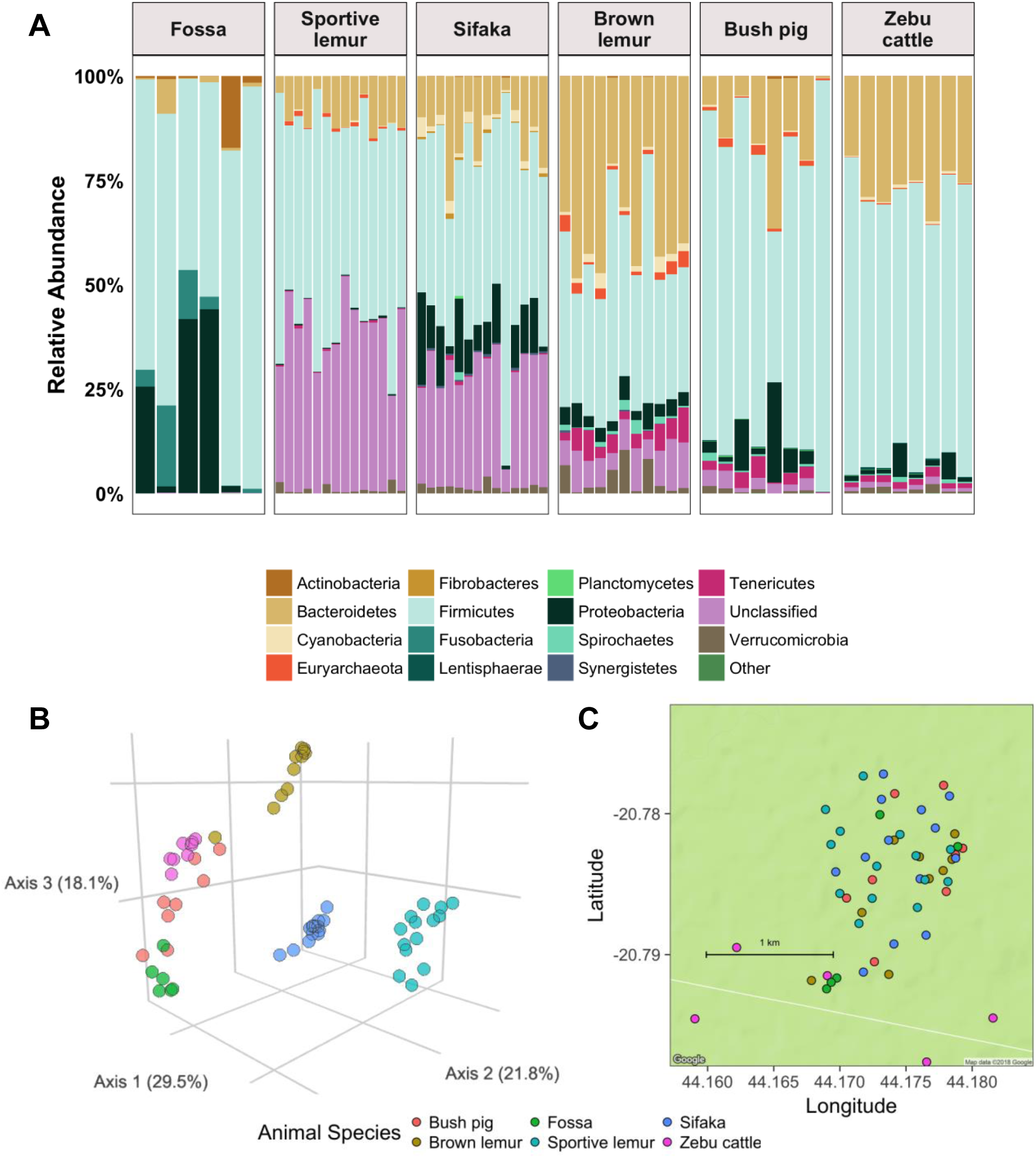
Sympatric mammal populations harbor distinct gut microbiotas. **A.** The relative abundances of the fifteen most abundant bacterial phyla in six Malagasy mammal species (fossa, *C. ferox*; red-tailed sportive lemur, *L. ruficaudatus*; Verreaux’s sifaka, *P. verreauxi*; red-fronted brown lemur, *E. rufifrons*; bushpig, *P. larvatus*; zebu cattle, *B. t. indicus*). The “Other” category represents low abundance (< 2%) phyla. **B.** Three-dimensional principal coordinates plot of normalized weighted Unifrac distances showing ecological distances among 61 microbiome samples from six mammal species. **C.** Fecal sample collection sites at Ankoatsifaka Research Station in Kirindy Mitea UNESCO Biosphere in western Madagascar.

### Host phylogeny and life history data

We estimated phylogenetic distances among mammal species using mitochondrial cytochrome b (cyt-b) gene sequences downloaded from NCBI and aligned using Muscle 3.8.31 (41). We evaluated models of molecular evolution (*phangorn* package) (42) and then reconstructed maximum likelihood phylogenetic trees based on the ‘TIM2+G’ model and 1000 bootstrap replications. Phylogenetic relationships among the distantly related species mirrored evolutionary times estimated from a time-calibrated ultrametric phylogenetic tree reconstructed with 5020 species (43), and those within the Strepsirrhine clade reflected divergence times estimated from a ultrametric Primate phylogenetic tree (44).

To quantify between-species differences in dietary intake, we used direct observations of feeding behavior for Verreaux’s sifaka at our research site (24) and, for the remaining five species for which we did not have direct observation data, primary literature specific to dry deciduous forests in western Madagascar (45–50). Following the EltonTraits database classification system (51), we defined ten dietary items (Invertebrates, Vertebrates excluding fishes, Reptiles/Amphibians, Carrion, Fish, Unknown Vertebrates, Fruits, Nectar/Pollen/Gums, Seeds/Nuts, Leaves/Grasses/Ground Vegetation), and each species was assigned a percentage for each dietary item. We then used Euclidean distances to compute a dietary distance matrix for the six species based on the proportions of dietary items consumed by each species. We broadly categorized zebu cattle as herbivorous (*i.e.*, feeding on plant material), sifaka and sportive lemurs as folivorous (*i.e.*, primarily feeding on plant photosynthetic material), brown lemurs as semi-frugivorous (*i.e.*, primarily feeding on fruit and seeds), bush pigs as omnivorous (*i.e.*, feeding on both plant and animal material), and fossa as carnivorous (*i.e.*, primarily consuming animal matter). We also used primary literature to categorize each mammal species by substrate use – ‘terrestrial’, ‘arboreal’, or ‘semi-terrestrial’– and gut physiology – ‘hindgut-fermenter’, ‘foregut-fermenter’, or ‘simple-gut’ (Table S2).

### Fecal sample DNA extraction and 16S amplicon sequencing

We extracted DNA from 100-200μg fecal pellets using a phenol chloroform bead-beating procedure (52). We quantified DNA samples with Picogreen® reagent (ThermoFisher) on a Qubit® fluorometer (ThermoFisher) and PCR amplified the V4 hypervariable region of the bacterial 16S ribosomal RNA gene using primers 515F and 806R (53). The resulting barcoded amplicons were pooled and paired-end sequenced in 2 × 151 mode using the Illumina MiSeq platform at Argonne National Laboratory (Lemont, IL).

### Sequence processing and taxonomic assignments

We de-multiplexed and quality filtered raw Illumina sequence reads using QIIME Version 1.8.0 (54). We paired forward and reverse reads using join_paired_ends.py and assigned reads to their respective samples based on their identifying barcode with split_libraries_fastq.py, allowing a minimum quality score of 20 and no errors in the barcode. We assigned reads attaining these quality standards to 97% Operational Taxonomic Units (*i.e.*, bacterial phylotypes) using the *uclust* algorithm and assigned taxonomic classifications with the RDP classifier (pick_open_reference_otus.py) based on their best match in the Greengenes database (13_5 release). After initial quality filtering, our dataset contained 4,556,629 processed paired-end sequence reads, averaging 74,699 reads per fecal sample. We removed singletons, OTUs designated as ‘unclassified’ at the kingdom level, and OTUs annotated as chloroplast or eukaryotic mitochondria to generate a usable OTU table of bacterial taxa. The resultant set of usable 97% OTUs contained a total of 3,747,987 reads (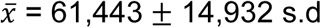. reads per sample, range: 27,632 — 88,491) and 98,278 unique 97% OTUs(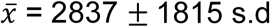. phylotypes per sample, range: 740 — 8513).

### Microbial diversity and composition among animal species

We conducted all statistical analyses using the statistical computing software R version 3.2.4 (55) and used the *ggplot2* (56) and *cowplot* (57) packages and Plotly (58) for visualization.

### Gut microbial richness

To test for differences in within-sample richness, diversity, and evenness among microbiomes, we generated 100 OTU tables rarefied to 35,384 reads (the smallest library size in the dataset) for each individual sample. After rarefaction, individual samples contained 610 to 6255 unique OTUs (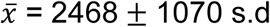. phylotypes per sample). We calculated mean rarefied richness (number of observed OTUs), Chao1 species richness, and Shannon’s diversity index for each host using the rarefied OTU tables. We used Kruskal-Wallis tests adjusted for multiple comparisons to evaluate whether bacterial taxa and evenness per individual differed across animal species.

### Sample clustering

We assessed similarity in gut microbial communities using only OTUs that were detected in at least two samples, totaling 3,508,792 reads (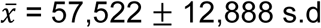. reads per sample, range: 26,847— 82,447) and 10,200 unique phylotypes. To account for differences in sequencing depth among samples and heteroscedasticity in OTU counts, we estimated sample-specific normalization factors using the *DESeq2* package (59) and then rescaled the OTU counts. After normalization, samples contained 203 — 1803 unique OTUs 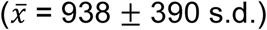. We conducted multivariate and community analyses using the *vegan* and *phyloseq* packages (60,61). We quantified inter-individual variation in gut microbial composition by calculating weighted Unifrac distances between samples and performed clustering of taxonomic profiles via partitioning of data around medoids (PAM) using the *cluster* and *fpc* packages (62). We used permutational multivariate analysis of variance (PERMANOVA) to assess differences in composition according to host species and substrate preference (1000 permutations) and a linear mixed-effect model to assess pairwise predictors of microbial similarity (methods provided in the supplementary information).

### Random forest classification

We determined the degree to which individual microbiome samples can be classified into their respective host species by implementing a random forest classifier (RFC) supervised learning algorithm (*randomForest* package) (63). We first eliminated OTUs with average relative abundances lower than 0.0001 and then used the remaining OTUs as predictors in the RFC model, with host species identity used as the category for RFC model distinguishability.

### Differentially abundant microbial taxa

We assessed differential abundance of bacterial phyla, families, and genera among host species using the nonparametric SAMseq approach (*samr* package) (64). This method uses repeated permutations for assessment of the false discovery rate (FDR). We limited analyses to bacterial and archaeal phyla that occurred at least 50 times, classifiable families that occurred at least 100 times, and classifiable genera that occurred at least 100 times in the dataset. We performed SAMseq analyses separately for each taxonomic level using 1000 permutations and 100 re-samplings and considered differential abundance significant if the FDR-adjusted *P* value was < 0.05.

### Mammalian microbiome meta-analysis

We combined our Malagasy samples with datasets extracted from primary literature on mammalian gut microbiota encompassing 47 mammal species across 13 orders (10,12). We assembled a closed-reference comparative data set with 101 samples (up to four samples per species chosen at random) and removed OTUs with less than 0.000001 relative abundance. Because we included 23 samples that were sequenced on the 454 platform (10), we rarefied the comparative dataset to 1100 sequences per sample to avoid biases in sequencing depth (12) and constructed an OTU table at the bacterial family level. We used non-metric multidimensional scaling to visualize Bray-Curtis dissimilarities among microbiome samples.

### Metagenomic predictions based on 16S rRNA sequencing data

We used PICRUSt (phylogenetic investigation of communities by reconstruction of unobserved states) v1.1.1 (65) to generate metagenomic functional profiles of lemur gut microbiomes based on patterns observed in 16S-based microbial community profiles. PICRUSt uses an extended ancestral-state algorithm to predict gene family abundances (*i.e.*, KEGG orthologs or pathways) from 16S data and a reference genome database. To visualize similarities in functional profiles among lemur species, we used redundancy analyses to construct principal component ordinations of KEGG orthologs and pathways. We created ordinations for the dataset as a whole (*i.e.*, all KEGG orthologs), as well as for individual KEGG sub-pathways related to carbohydrate and essential amino acid metabolism. We implemented an RFC model to determine the degree to which individual predicted metagenomes can be classified into their respective host species.

### Microbiome composition and host phylogeny

We used a parsimony approach (8) to determine whether similarities in gut microbiome composition are congruent with the evolutionary relationships among mammalian host species. We used phylogenetic methods to infer the pattern of hierarchical similarity among hosts based on the frequencies and abundances of taxonomically assigned OTUs in their gut microbiomes. Because unique OTUs cannot provide information about the evolutionary relatedness among samples, we limited our phylogenetic analysis to OTUs detected in two or more samples, which included 12,967 OTUs. We first created a character matrix comprised of these phylogenetically informative OTUs, with each character corresponding to an OTU whose absolute abundance was normalized by coding into one of six states reflecting log-unit differences in occurrence, with ‘1’ corresponding to an OTU’s absence in a given sample. Characters were ordered, such that transitions between distant states (*i.e.*, samples having very divergent frequencies of a particular OTU) were costlier than between similar states. We then subjected the six-state character matrix to a heuristic maximum parsimony tree search (*phangorn* package) (42), with 1000 pseudo-replicates used to assess bootstrap support, and recovered a single maximum parsimony tree (p-score = 35,672). We used the normalized Robinson-Foulds score and Mantel tests to assess congruence between host and gut microbiome topologies (methods provided in the supplementary information).

## Results

### Microbiome composition

The gut microbial communities of the six mammalian host species encompassed one archaeal and 28 bacterial phyla, of which three (Firmicutes, Proteobacteria, and Bacteroidetes) were present in all samples and together constituted 48-99% of the reads identified in each individual (Fig. 1A). The microbial phyla, families, and genera associated with each animal species are detailed in the supplementary information (Fig. S2, Table S3). Bushpig and cattle microbiomes were enriched with Planctomycetes and Lentisphaerae whereas fossa microbiomes were deficient in Bacteroidetes and showed higher abundances of microbial taxa related to Actinobacteria and Fusobacteria (SAMseq analysis, FDR-adjusted *P* < 0.0001; Table S3). The microbiomes of the two folivorous lemur species, sifaka and sportive lemurs, included higher proportions of unclassified bacterial phyla (*P* < 0.0001). Sifaka and brown lemurs, the most closely related mammal species in our study, tended to share bacterial phylotypes belonging to Cyanobacteria and Verrucomicrobia (*P* < 0.0001; Table S3). Lastly, microbial lineages related to Tenericutes and Spirochaetes that were comparatively abundant in brown lemurs were also enriched in cattle and bush pig samples (*P* < 0.0001; Table S3). Although we did not have data to show the presence of specific pathogens in our samples, several bacteria genera that include opportunistic pathogens, such as *Campylobacter, Clostridium*, and *Streptococcus*, were differentially abundant across mammal species (Fig. S2, Table S3).

In accordance with (13), we found that differences in gut biodiversity among mammal species reflected digestive physiology, diet, and host taxonomy. Cattle (herbivorous foregut fermenting artiodactyls) and bush pigs (omnivorous artiodactyls with simple gut morphology) harbored the most diverse microbiota (Kruskal-Wallis, FDR-adjusted *P* < 0.001; Fig. S3), whereas the three lemur species, two of which are hindgut fermenters (66,67), exhibited moderate biodiversity (Fig. S3). Fossa, the only carnivorous species in our study, had the fewest unique taxa and the lowest microbial species evenness (*P* < 0.001; Fig. S3).

### Microbiome diversity across sympatric host species

Microbiome samples formed five to six discrete clusters (PAM clustering, Fig. S4) according to host species identity (PERMANOVA R^2^ = 0.68, *P* = 0.001; RFC 100% cross-validation accuracy), with sifaka and sportive lemur microbiomes clustered independently and brown lemur microbiomes grouped more closely with those of cattle, bush pigs, and fossa (PCoA, Fig. 1B). The bacterial communities of the three lemur species were taxonomically similar at the phyla level (Fig. 1A), but distinct when classified by 97% phylotype similarity (Fig. 1B). Microbiome distances between species correlated significantly with substrate use (*i.e.*, terrestriality versus arboreality) (PERMANOVA, R^2^ = 0.16, *P* = 0.001; GLMM, *P* < 2 × 10^−4^, Table 1). Terrestrial and semi-terrestrial species exhibited similar microbial taxonomic structure and abundances (*i.e.*, closer weighted Unifrac distances) despite having divergent diets and gut physiologies. In contrast, the microbiomes of the arboreal sifaka and sportive lemurs were highly divergent, despite their shared substrate preference and leaf-based diets (Fig. 1B).

**Table 1:** Pairwise predictors of weighted Unifrac distance among six sympatric mammal species. Posterior mean, 95% credible interval (95% CIs), and *P*-value based on Markov chain Monte Carlo sampling for fixed-effect parameters. Baseline pairwise terrestriality (“one or both species are not terrestrial”) is not shown. Individual sample identity within each pair was included as a random effect. Bolded relationships are significant at pMCMC < 0.05.

| Parameter | Mean | 95% CI | pMCMC | Interpretation |
| --- | --- | --- | --- | --- |
| Intercept | 0.21 | (0.19, 0.23) | **$< 2 \times 10^{-4}$** |  |
| Dietary distance | $1.61 \times 10^{-4}$ | $(6.06 \times 10^{-5}, 2.56 \times 10^{-4})$ | **0.0009** | Host species with divergent diets have more dissimilar microbiota |
| Both terrestrial | -0.03 | (-0.04, -0.02) | **$< 2 \times 10^{-4}$** | Host species that both spend time on the ground have less dissimilar microbiota |
| Phylogenetic distance | 0.3 | (0.29, 0.31) | **$< 2 \times 10^{-4}$** | Host species that are more evolutionary distant have more dissimilar microbiota |
| Geographic distance | $-2.36 \times 10^{-3}$ | $(-6.39 \times 10^{-3}, 1.78 \times 10^{-3})$ | 0.23 | No significant correlation |

### Metagenomic functional analyses of lemur microbiomes

The gut microbiomes of the three lemur species had significantly distinct metabolic functionality (RFC 92.3% cross-validation accuracy, Fig. S6). Given the phylogenetic proximity of these species, we hypothesized that these functional differences stem from diet intake rather than host evolutionary divergence. Sportive lemur microbiomes were enriched for both sugar metabolism and plant fiber degradation, consistent with a combined frugivorous and folivorous diet (Fig. 2). In contrast, brown lemur microbiomes were enriched for only sugar metabolism, with elevated levels of enzymes for metabolizing fructose, mannose, and sucrose, consistent with a fruit-based diet (Fig. 2). Sifaka microbiomes prioritized fiber degradation pathways, with enzymes for butanoate, propanoate, and glyoxylate metabolism, consistent with a leaf-based diet (Fig. 2). Pathways related to essential amino acid metabolism also distinguished the gut microbiotas of the lemur species and reflect disparities in protein content between fruit- and leaf-based diets (Fig. S6). Fruit pulp tends to be low in protein content, whereas leaves contain considerable amounts of rubisco, a protein involved in photosynthesis (68). Hence, pathways for the biosynthesis of lysine and branched-chain amino acids (valine, leucine, isoleucine) were elevated in brown lemurs and sportive lemurs, whereas the catabolic reactions to break down these amino acids were enriched in sifaka (Fig. S6).

**Figure 2.**
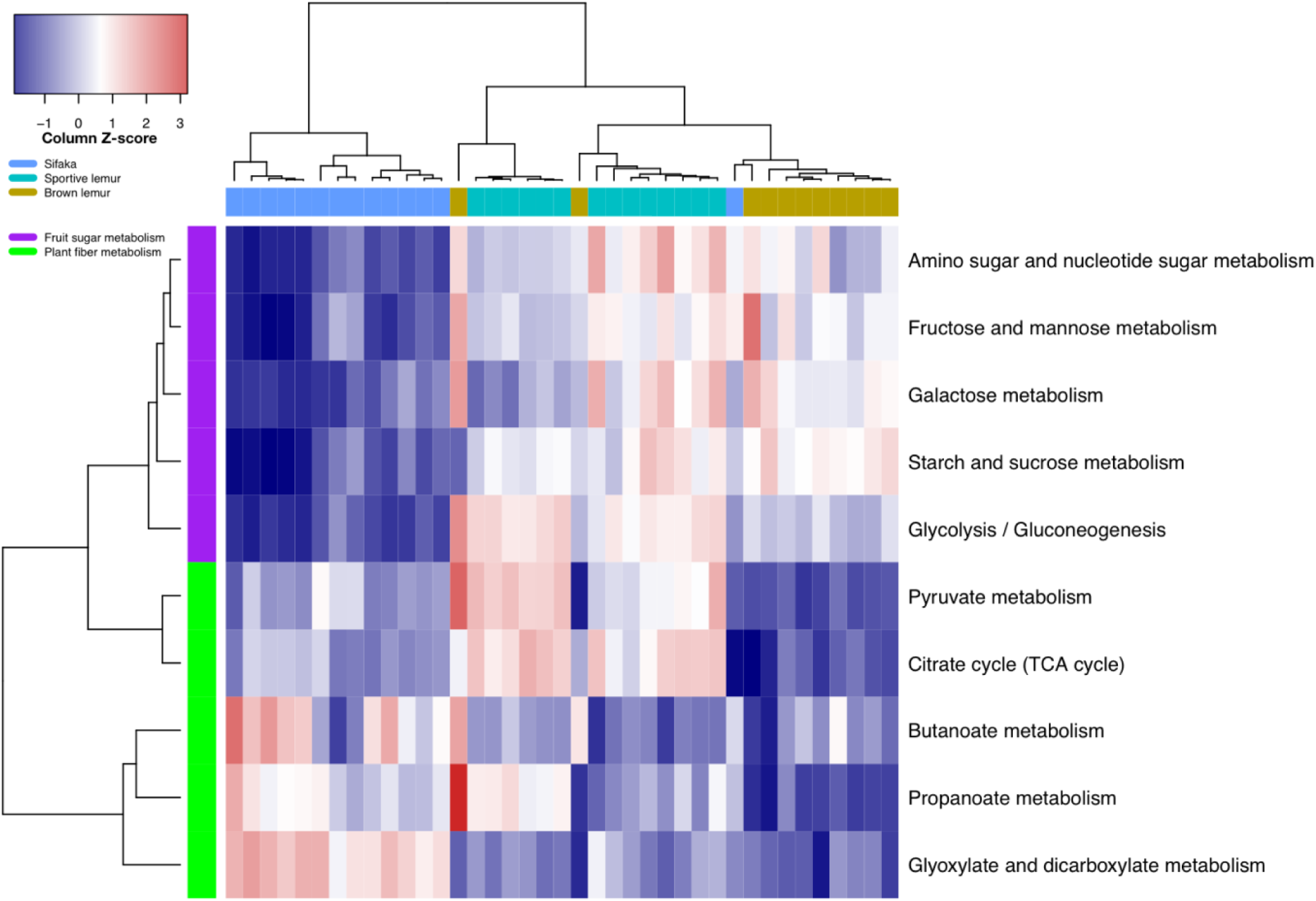
Heatmap of predicted carbohydrate metabolism in brown lemur, Verreaux’s sifaka, and sportive lemur microbiomes. The transformed relative abundances of the ten most abundant carbohydrate KEGG sub-pathways are shown, with the white color representing the relative abundance of pathways having the column average, blue tones indicating relative abundances less than the column average, and red tones indicating relative abundances greater than the column average. Brown lemur microbiomes are enriched for pathways associated with a frugivorous diet (purple rows), sifaka for those associated with a folivorous diet (green rows), and sportive lemurs for both frugivorous and folivorous pathways. Metagenomic functional profiles were inferred using PICRUSt.

### Microbiome phylogenies mirror host phylogenies

We reconstructed gut microbiota phylogenies based on the frequencies and abundances of microbial phylotypes among microbiome samples (Figs. 3 and S7). The microbiota tree branching mirrored the host phylogeny for distantly related species (nRF = 0.33, *P* = 0.0001), but not for the more closely related sportive lemurs, brown lemurs, and sifaka (Figs. 3 and S7). Specifically, the sportive lemur and brown lemur microbiomes were more closely related in the microbiota tree than in the host tree. This finding is consistent with the metagenomic functional patterns and suggests that diet may play a greater role than evolutionary history in structuring microbial communities among closely related sympatric primate species. Despite incomplete concordance between the host mtDNA and gut microbiota phylogenies, host mitochondrial genetic distance was a significant predictor of weighted Unifrac distance among microbial communities (Mantel, r = 0.81, *P* = 0.001), even after controlling for dietary distance among host species (partial Mantel, r = 0.74, *P* = 0.001) and geographic distance among fecal sample collection sites (partial Mantel, r = 0.82, *P* = 0.001).

**Figure 3.**
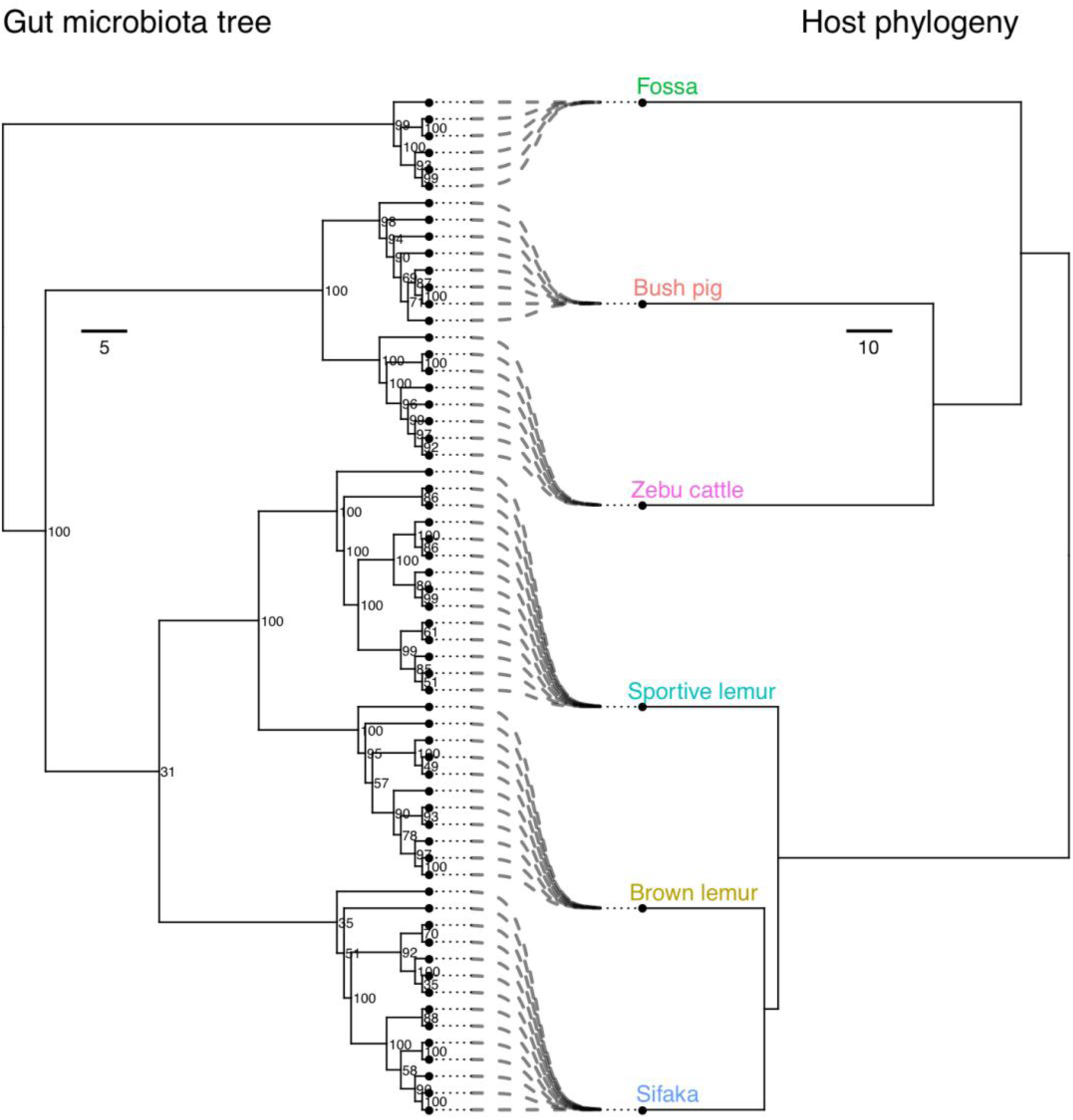
Comparison between gut microbiota tree and host phylogeny relationships. The gut microbiota tree reflects the same branching order as the host phylogeny for distantly related host species, but gut microbial similarity among sportive lemurs, brown lemurs, and sifaka does not match these hosts’ established evolutionary relationships. The maximum parsimony gut microbiota tree (left) is based on the frequencies and abundances of 97% OTUs in 61 microbiome samples from six mammal species. The maximum likelihood host phylogeny (right) was inferred using mitochondrial cytochrome b (cyt-b) gene sequences.

We further compared these Malagasy mammalian gut microbiomes to those of 46 other mammal species among 13 host orders (10,12). Bush pigs and cattle clustered with other members of Artiodactyla, lemurs with other primates, and fossa microbiomes with other carnivores (Figs. 4 and S8). In a meta-analysis, both host order and diet contributed significantly to variation in gut microbiome composition among the 52 animal species (PERMANOVA: host order: R^2^ = 0.3, *P* = 0.001; diet: R^2^ = 0.24, *P* = 0.001; Figs. 4 and S8).

**Figure 4.**
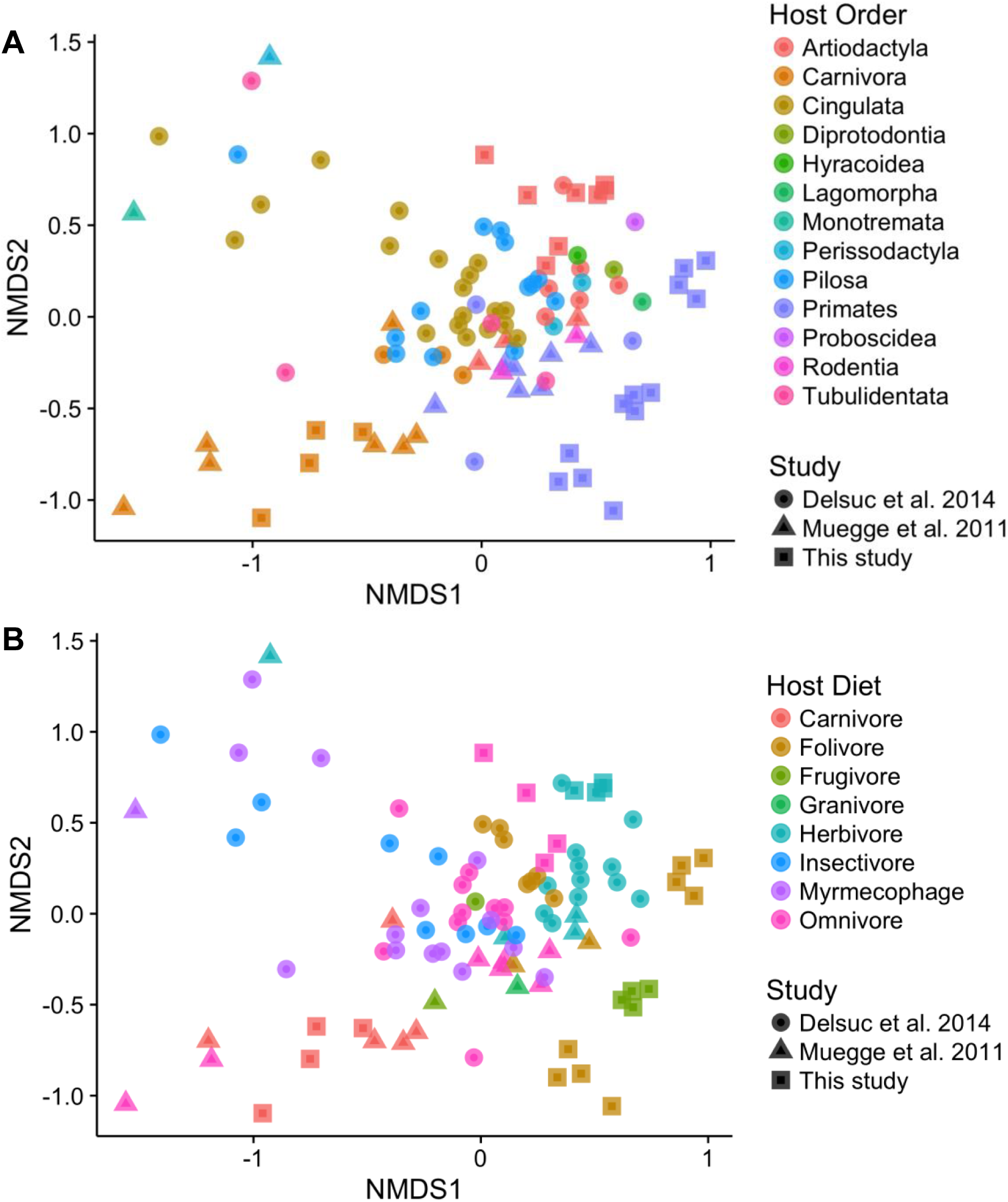
Malagasy mammal microbiomes cluster according to host order (A) and diet (B). Non-metric multidimensional scaling ordinations of family-level Bray-Curtis dissimilarities among mammalian gut microbiome samples from 52 host species across 13 orders. Samples from this study were combined with those from two other comparative studies of captive and wild animal microbiota (10,12).

## Discussion

We investigated the ecological and evolutionary determinants of gut microbiome composition among six sympatric mammal species co-residing within a 1-km^2^ area in western Madagascar. As found previously (10–12,14), variation in taxonomic composition among wild Malagasy mammals was correlated with both host evolutionary history and diet, depending on the relatedness of the species. Moreover, shared terrestriality, but not geographic distance between sampling sites, predicted microbiome similarity among distantly related hosts with differing diets and digestive physiologies. This suggests that ground dwelling promotes the indirect horizontal transmission of commensal gut bacteria among sympatric wild mammals.

Gut bacterial diversity reflected a combination of digestive physiology, diet, and host phylogeny. The three feeding strategies represented in our study – *carnivory* and two types of herbivory, *folivory* and *frugivory –* range from foods that are energetically costly to obtain but easy to digest (*i.e.*, animal matter), to those that are limited in quantity but fairly easy to digest (*i.e.*, fruits and seeds), to those that are ubiquitous but difficult to digest (*i.e.*, leaves) (69). Carnivorous fossa and frugivorous brown lemurs require only short, simple guts because the protein and fat in animal matter and the short-chain sugars in fruit are easily assimilated using enzymes produced by the animals themselves (69,70). In contrast, the herbivorous and folivorous mammals (zebu cattle, sportive lemurs, and sifaka) have complex gastrointestinal tracts with lengthened digesta retention times and diverse communities of mutualistic microorganisms to facilitate fiber digestion (71,72). In agreement with a prior survey of 59 mammalian host species (11), we found that bush pigs and cattle, both members of the foregut-fermenting Artiodactyla order, exhibited the greatest bacterial diversity, followed by the three lemur species with intermediate diversity, and finally carnivorous fossa with lowest bacterial richness. Given that sifaka and sportive lemurs, both folivorous hindgut fermenters (73,74), did not harbor significantly greater bacterial richness than brown lemurs, which are frugivorous (45), we conclude that microbiome diversity within the lemur clade is shaped more by host phylogeny than digestive physiology.

The gut microbiomes of the six mammal species were distinguishable with 100% accuracy, and host phylogeny was the strongest predictor of compositional similarity, even when accounting for geographic proximity of sampling sites, substrate preference (*i.e.*, ground versus tree), and dietary overlap among host species. With the exception of the lemur clade, the microbiomes of sympatric Malagasy mammal populations illustrate “phylosymbiosis,” an eco-evolutionary pattern in which the evolutionary diversification of hosts correlates with ecological changes in their microbiota (18,75). Congruence between host and microbial phylogenies can arise without the coevolution of entire microbial communities from a last common ancestor or co-speciation events (9,18). Thus, phylosymbiosis does not presuppose that all resident phylotypes in microbiomes are stable or vertically transmitted between generations (9,18). For example, closely related hosts with comparable behavioral traits may acquire similar bacteria from their environment or directly from other hosts (9,76). In mammals, phylosymbiosis has been observed in large-scale comparative studies (9), among closely related species in controlled environments (18), and in wild populations of bats (16) and apes (8,77,78).

The branching order of lemur microbiome samples did not reflect the established evolutionary relationships among brown lemurs, sportive lemurs, and sifaka. Sifaka and sportive lemur microbiomes formed separate, distant clusters while brown lemur microbiomes grouped more closely with those of cattle, bush pigs, and fossa. This finding was surprising, given that sifaka and sportive lemurs are closely related species with overlapping habitat use and diet, whereas the four semi-terrestrial and terrestrial species are distantly related with differing diets and digestive physiologies. A recent study found that microbiome composition and host phylogeny are strongly associated for recently diverged captive mammals (9). In contrast, it seems that diet and substrate preference are more influential than host phylogeny in shaping gut microbial composition among sympatric wild lemur species.

The climate of western Madagascar is highly seasonal. Many Malagasy lemur species modify their diet composition and foraging strategies during the dry season as temperatures fluctuate and high quality food resources and water become scarce (45,79). The *Eulemur* clade are flexible frugivores (45), with high levels of folivory recorded in western populations during the dry season (80,81). *Propithecus* and *Lepilemur* are both considered folivorous (46,73,82,83), consuming leaves throughout the year (45,46,84) and fruit when available during the rainy season (45,46,85,86). We found that the predicted functional composition of lemur microbiomes reflected differences in carbohydrate and amino acid utilization, with red-fronted brown lemurs (*Eulemur rufifrons*) displaying frugivore dietary profiles, Verreaux’s sifaka (*Propithecus verreauxi*) folivore profiles, and red-tailed sportive lemurs (*Lepilemur ruficaudatus*) intermediate frugivore-folivore dietary profiles. As found for captive lemurs (87), both sportive lemur and sifaka microbiomes were enriched for folivorous pathways associated with increased plant fiber degradation, such as propanoate and butanoate metabolism (88). Along with brown lemur microbiomes, sportive lemur microbiomes were also elevated for frugivorous pathways related to sugar metabolism and amino acid biosynthesis. Thus, frugivory may explain the gut microbial similarity between these two species, despite the more recent evolutionary divergence between sifaka and brown lemurs. Seasonal frugivory observed in captive sportive lemurs supports this claim (46). However, there have been insufficient dietary studies of *Lepilemur* spp. in dry deciduous forests to definitively compare the fruit consumption of wild sportive lemurs and sifaka (46,86).

The striking discordance between sifaka and sportive lemur microbiomes may also stem from differences in host body size, activity patterns, and digestive strategies. Sifaka are diurnal with a body mass of 2.5—4 kg (89), while sportive lemurs are remarkably small-bodied (< 1kg) for a folivorous species (90). Large body size is considered a morphological adaptation to folivory, providing increased gastrointestinal surface area and retention time for nutrient absorption (69). Caecotrophy—re-ingestion of feces—has been observed in one sportive lemur species (*L. mustelinus)* and may be an adaptation to increase nitrogen utilization (82). However, caecotrophy has not been observed in our study species (*L. ruficaudatus*) or other sportive lemurs (86,90,91). Instead, they may manage their folivorous diet and small body size by conserving energy; they have one of the lowest basal metabolic rates among folivorous mammals (92) and long nighttime resting periods (82,90). Although sifaka and sportive lemurs are both hindgut fermenters, their digestive strategies lie at opposite ends of the continuum observed for primates. Sifaka employ an “efficiency” strategy, characterized by low intake, long mean retention time, and high fiber digestibility (93), while sportive lemurs use an “intake” strategy, with high intake, short mean retention time, and low fiber digestibility (70,84). Thus, the gut microbiota of sifaka may be better suited for degrading structural carbohydrates and detoxifying plant secondary compounds.

Finally, shared terrestriality was a significant predictor of microbial similarity, with the microbiomes of semi-terrestrial brown lemurs and fossa clustered closely to those of terrestrial bush pigs and cattle. Although a prior meta-analysis of primate parasite diversity failed to find a substrate effect (94), other studies have reported lower parasite prevalence in arboreal primates compared to sympatric terrestrial primates (95–97). These studies did not, however, account for disparities in home range size, which may influence exposure to microorganisms (98). Thus, terrestrial and semi-terrestrial animals may have greater exposure to fecal-orally transmitted microorganisms and soil-borne parasites compared to their arboreal counterparts (97,98). The arboreal lifestyles and comparatively smaller home ranges of sifaka and sportive lemurs may limit incidental contact with enteric bacteria in the environment (95,98), and thus compound the effects of their unique diets and physiologies in driving the divergence of their gut microbial communities.

Although we focused on commensal and mutualistic gut microbiota, our findings may also apply to pathogenic bacteria that exploit similar molecular mechanisms to colonize hosts (99). We speculate that microbiome overlap among distantly related terrestrial species is driven by exposure to heterospecific fecal material on the ground. Terrestrial animals may therefore be more vulnerable to cross-species spillover of enteric bacteria compared to arboreal animals (95,97,98). A recent study (29) reported gut bacterial similarities within predator-prey host-species pairs in North America, suggesting that mammalian food chains may serve as transmission routes for gut bacteria. In contrast, sifaka and sportive lemurs constitute a high proportion of the fossa diet in western deciduous forests (47,48), and we found that the microbiomes of these three species were markedly distinct.

## Conclusions and Future Directions

Our comparative survey of mammalian gut microbial communities is one of very few to assay cohabiting wildlife populations and is unique in its small geographic extent. By eliminating the potentially confounding effects of geography and captivity, we are able to elucidate the relative contributions of diet, behavior, and evolutionary history on gut microbial diversity in understudied Malagasy mammalian species. In contrast to prior microbiome surveys of geographically distinct or captive mammal populations (8–12,17), we find that the gut microbiomes of distantly related species reflect their hosts’ evolutionary relationships rather than their dietary classifications. However, for closely related primate species, diet and substrate use, rather than phylogeny, seem to be the driving factors. Epidemiological studies in wildlife have elucidated cross-species transmission of pathogens (94,100), but little is known about the transfer of commensal microbes among co-occurring wild animal populations (28,29). More comprehensive comparative studies of wild mammal microbiomes are needed to fully resolve the dynamic dietary, social, and environmental factors that constrain gut microbiome ecology and evolution. Such insights will advance our understanding of host-microbe coevolution and the evolution of mammalian dietary flexibility and diversification and improve strategies for wildlife conservation and captive management.

## Acknowledgements

We thank the Madagascar government, CAFF/CORE, and Madagascar National Parks for permission to conduct this research, the University of Antananarivo, MICET, and the Ankoatsifaka Research Station staff for facilitating our research, and Elvis Rakatomalala for his help with fecal sample collection. We thank Anthony Di Fiore, Maryjka Blaszczyk, and Kelsey Ellis for their helpful comments on the manuscript. This work was supported by the National Institutes of Health/MIDAS, the National Science Foundation, the BEACON Center for the Study of Evolution in Action, the International Primatological Society, the American Society of Primatologists, and The University of Texas at Austin.

## Data availability

The data that support the findings of this study will be made publicly available in the Sequence Read Archive at the time of publication acceptance.

## Authors’ contributions

ACP, RJL, and LAM designed the research; ACP collected fecal samples, analyzed the data, and wrote the manuscript. All authors revised drafts of the manuscript and gave final approval for publication.

